# Cultivation and physiological characterization of a desert-derived *Halospirulina* isolate

**DOI:** 10.64898/2026.05.27.728284

**Authors:** Bárbara Bastos de Freitas, Mirian dos Santos Mendes, Mark Pampuch, Mauricio Lopez Portillo Masson, Kyle J. Lauersen

## Abstract

Here, we describe a filamentous *Halospirulina* isolate (*Halospirulina saudiensis*) obtained from water-clay microhabitat the Empty Quarter desert, الربع الخالي (ar-Rubʿ al-Khālī), Saudi Arabia which grows in saline conditions. We present its fully sequenced genome, the first for the genus, and characterize its growth dynamics as well as biochemical composition under a range of cultivation conditions. Protein, carbohydrate, lipid, and phycocyanin content varied with cultivation regime but were largely stable. *H. saudiensis* reached biomass concentrations of up to 9.83 g L^-1^ at pH 7, 35 °C and continuous 325 µmol photons m^-2^ s^-1^. Variable climate simulations in lab-scale photobioreactors revealed preference for warmer season cultivation under modeled outdoor conditions. Carotenoid analysis revealed a pigment profile enriched in canthaxanthin and other ketocarotenoids, distinguishing it from industrial *Limnospira* and positioning its value for neutraceuticals and feed additives. Genome analysis identified a carotene ketolase (*crtO*) homolog consistent with other cyanobacteria that accumulate ketocarotenoids. Phycocyanin content was heavily dependent on culture health and varied with cultivation pH, irradiance, reaching maximum values of 67.3 ± 0.8 mg gDW^-1^ (6.73 %). Extracted phycocyanin showed marginal thermal stability compared to that from *L. platensis*. The findings suggest that *H. saudiensis* could be a promising source of biomass, ketocarotenoids, and natural pigments, cultivated in saline conditions with elevated temperature and irradiance.

**Graphical Abstract:** 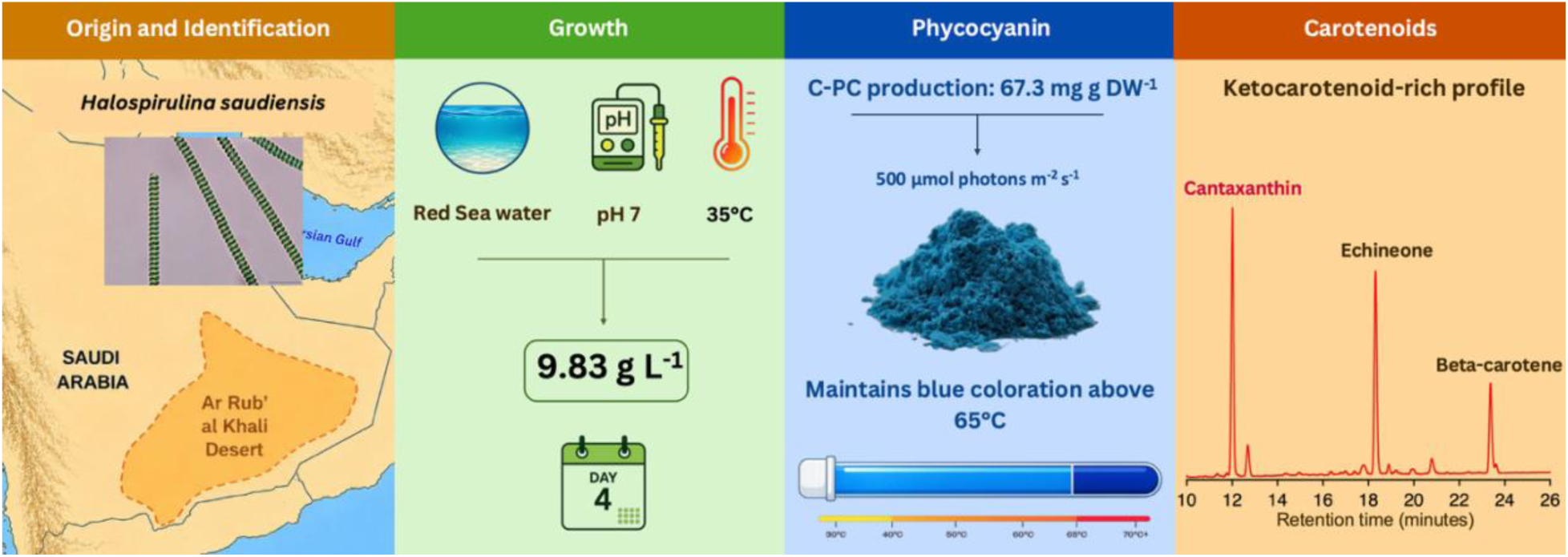

**Highlights:**

- *Halospirulina saudiensis* sp. nov. isolated from Empty Quarter
- First genome-resolved characterization of a *Halospirulina* strain
- Reached 9.83 g L^-1^ in Red Sea salinity conditions
- Accumulates canthaxanthin as major carotenoid
- Phycocyanin slightly thermotolerant

## 1 Introduction

Microalgae and cyanobacteria are biological resources that can be cultivated for the production of food, feed, pigments, and other high-value biomolecules (Hosseini et al., 2026). Among these organisms, commercial ‘Spirulina’ cultivation is primarily based on strains within the genus *Limnospira* (formerly *Arthrospira*), which are widely used for nutritional supplements and the production of blue, water soluble phycocyanin (C-PC) (Demirel & Sukatar, 2019; Kobbin et al., 2025). *Limnospira* are currently among the most widely cultivated microalgae at commercial scale, with production predominantly carried out in open raceway ponds and reaching tens of thousands of tons of biomass annually (González-Portela et al., 2024; Lopes et al., 2026). The global market has been estimated at approximately USD 619 million and is projected to approach USD 1 billion by the end of the decade (Hosseini et al., 2026). This demand is supported by the biomass composition of *Limnospira*, particularly its high protein content and production of C-phycocyanin, enabling applications in nutraceuticals, functional foods, cosmetics, and aquaculture (Hosseini et al., 2026; Vasquez Guevara et al., 2025). However, commercial cultivation remains dependent on large volumes of water and relatively controlled cultivation conditions to maintain productivity and reduce contamination (Li et al., 2024; Yi et al., 2025). These constraints have increased interest in alternative cyanobacterial strains adapted to cultivation under less competitive environmental conditions.

The taxonomy of filamentous cyanobacteria historically classified as *Spirulina* has undergone substantial revision following the application of molecular phylogenetic approaches (Komárek et al., 2014; Nowicka-Krawczyk et al., 2019; Nübel et al., 2000). Sequence-based analyses demonstrated that morphologically similar spiraled cyanobacteria belong to multiple phylogenetic lineages, resulting in the recognition of several distinct genera within the former *Spirulina* complex (Komárek et al., 2014; Nübel et al., 2000). One of these lineages corresponds to the genus *Halospirulina*, a group of filamentous cyanobacteria commonly associated with saline and alkaline environments (Szubert et al., 2021; Waditee-Sirisattha & Kageyama, 2023). Despite their ecological distribution, comparatively few studies have investigated the cultivation performance, biomass composition, or biotechnological potential of *Halospirulina* strains relative to commercially exploited *Limnospira* species.

Environments characterized by elevated salinity, intense irradiance, and strong temperature fluctuations are recognized as important sources of phototrophic microorganisms with physiological traits relevant for biotechnology (Hosseini et al., 2026; Kumawat et al., 2025; Markou et al., 2023). Desert ecosystems, in particular, expose microorganisms to high irradiance, elevated and diurnally dynamic temperatures, and prolonged water limitation, conditions that may select for strains with resilient and biochemically interesting metabolism. The ‘Empty Quarter’, الخالي الربع (ar-Rubʿ al-Khālī), desert in Saudi Arabia represents a highly arid environment and remains comparatively underexplored as a source of cyanobacterial diversity (Edgell, 2006).

Here, samples collected from a water and clay microhabitat beside a desert road embankment, associated with a shallow abandoned anthropogenic evaporation area, led to the isolation of a previously undescribed *Halospirulina* strain. The isolate was characterized here for its growth performance, biomass composition, unusual carotenoid profile, and phycocyanin production under different pH, temperature, irradiance, as well as in simulated seasonal cultivation conditions representative of outdoor cultivation on the mid Red Sea coast of Saudi Arabia.

## 2 Materials and methods

### 2.1 Field sampling, enrichment and strain isolation

Field sampling was conducted during Winter 2024 in the Empty Quarter desert, Saudi Arabia at 20.656259536743 N, 54.63586807251 E. The site consisted of a shallow saline water body, potentially associated with an evaporation pond formed during historical road construction activities in the region. Water-clay samples were trasferred to sterile 175 mL vent-cap culture flasks for transport to the laboratory for enrichment and isolation.

For enrichment, approximately 1 g of water-clay sample was suspended in 50 mL of sterile culture medium in vent-cap culture flasks. BG-11 medium (Rippka et al., 1979) prepared with freshwater, BG-11 or F/2 medium (Guillard & Ryther, 1962) prepared with ultrafiltered and autoclaved Red Sea seawater. The isolate described in this study was obtained from enrichment culture grown in BG-11 made with Red Sea water at 42 °C. Sodium nitrate (NaNO_3_) was supplied as the nitrogen source in all media. Cultures were maintained at 25 ± 3 °C or 42 ± 3 °C, under continuous illumination of approximately 150 µmol photons m^-2^ s^-1^, with gentle agitation. During enrichment period (14-20 d) and isolation, cultures were periodically examined by light microscopy to monitor cyanobacterial growth and evaluate culture purity. Serial dilution was used to establish algal monocultures. Morphological observations were performed using a BX53 upright microscope (Olympus, Tokyo, Japan) equipped with a CMOS color camera (SC50). Following isolation, the strain was maintained in the KAUST KSA Living Library maintained by the Sustainable and Synthetic Biotechnology Group at KAUST (https://ssb.kaust.edu.sa/KSALivingLibrary).

### 2.2 DNA extraction, sequencing, and strain identification

Cell cultures were harvested by centrifugation at 3000 x *g* for 20 min and pellets washed with 1X phosphate-buffered saline (PBS). High-molecular weight DNA (HMWDNA) was extracted following the QIAGEN Genomic DNA Handbook protocols for cultured cells using Genomic-tip 500/G columns (QIAGEN, Cat. no. 10262). Sequencing libraries were prepared using the PacBio HiFi plex prep kit 96 (Product Number: 103-381-300) and sequenced on the PacBio Revio system. Genome assembly was performed using the nf-core/bacass pipeline (v2.5.0, Daniel et al., 2025) with dragonflye assembler (v1.2.1). Predicted rRNA sequences were extracted from the genome assembly contigs with barrnap (v0.9, Seemann, 2018) using the ‘--kingdom bac’ parameter and subsequently searched against the NCBI nucleotide database with BLAST. A circularized 5 Mb contig containing the isolate’s-affiliated rRNA sequences was selected as the final genome assembly used in this study. Genome annotation was performed using BAKTA (v1.6.x) (Schwengers et al., 2021) with the BAKTA database v6.0 (Schwengers, 2025).

Phylogenetic analysis was performed by extracting the 16S rRNA sequences from the isolate’s genome using the bacterial small subunit ribosomal RNA covariance model obtained from Rfam database (version 15.1; accession: RF00177) (Ontiveros-Palacios et al., 2025), using the cmscan software (with default parameters) from the Infernal toolkit (v1.1.5) (Nawrocki and Eddy, 2013). The phylogenetic tree was generated by first extracting all *Spirulina*, *Halospirulina*, and *Arthrospira* 16S rRNA sequence alignments from the SILVA SSU database (release 138.2) (Chuvochina et al., 2026). Strains classified as *Limnospira* were not present in this version of the SILVA and, therefore, could not be included. In addition, SSU alignment data from *Nostoc* sp. BDU-40302 was also extracted to be used as an outgroup for phylogenetic tree construction. The first of two 16S rRNA sequences extracted from the isolate’s genome was used to construct the phylogenetic tree. This sequence was added to the existing SILVA multiple sequence alignment using MAFFT (v7.505) (using the ‘--add’, ‘--anysymbol’, and ‘--keeplength’ parameters). IQ-TREE3 (Wong et al., 2025) was used to generate the phylogenetic tree on the multiple sequence alignment, using automatic model selection, 1000 replicates of the SH-aLRT test, and 1000 replicates of non-parameteric bootstrapping (parameters ‘-m MFP -alrt 1000 -B 1000’). Tree visualization was done using iTOL (Letunic & Bork, 2007)(Letunic & Bork, 2007).

For phylogenomics analysis, all available genomes from the family Spirulinaceae and the genera *Arthrospira* and *Limnospira* were retrieved from the Genome Taxonomy Database (GTDB r226.0) (Parks and Hugenholtz, 2026). Entries that had discrepancies between the NCBI and GTDB taxonomic classifications were subsequently omitted from the phylogenomic analysis, except in the cases where *Arthrospira* strains were classified as *Limnospira* strains, as these two genera have historically had intertwining classifications (Sinetova et al., 2024). Entries whose genomes failed GTDB genome quality control check were also omitted from the analysis. A representative from the Nostocaceae family (*Nostoc commune* HK-02; GCA_003990685.1) was chosen as an outgroup. The GTDB-Tk (Chaumeil et al., 2022) de_novo_wf was performed to extract marker genes from these genomes, perform multiple sequence alignments, and concatenate the results. IQTREE3 was used to generate the phylogenomic tree on the multiple sequence alignment generated by the GTDB-Tk workflow, using the same parameters as in the phylogenetic tree described above (‘-m MFP -alrt 1000 -B 1000’). Tree visualization was done using iTOL. Average Nucleotide Identity (ANI) was performed using anvi’o v0.9 (Eren et al., 2021) with the anvi-compute-genome-similarity workflow (with the ‘ --program pyANÌ flag set) and pyANI using the ANIb module. The minimum alignment fraction was set to 0.5. Protein sequences listed in Supplementary material 1 were retrieved from UniProt and used as queries for homology screening. Homologs were identified by performing tBLASTn searches against the target genome using default parameters.

### 2.3 Cultivation conditions

Cultivations were conducted in biological duplicates in Algem photobioreactors (Algenuity, UK) using BG-11 medium (Rippka, 1979) prepared with natural Red Sea seawater. Unless otherwise stated, cultivations were performed for 7 d in duplicate using 400 mL working volumes with agitation at 120 rpm. A gas mixture containing 3% CO_2_ in air was periodically supplied at 25 mL min^-1^ for pH control. Biomass accumulation was monitored through dry weight measurements. For each replicate, 9 mL of culture was collected as three 3 mL aliquots, centrifuged into pre-weighed glass tubes, and dried at 100 °C for 24 h before weighing. To test the effect of pH on culture growth, cells were grown at 35 °C and 40 °C under pH 7, 8, and 9 controlled by CO_2_ sparging with 325 µmol photons m^-2^ s^-1^. Samples collected at the end of cultivation were used for biomass composition and phycocyanin analyses. Irradiance was tested in cultures maintained at pH 7 and 35 °C using illumination at 200, 325, and 500 µmol photons m^-2^ s^-1^. Biomass samples collected on d4-7 were used for macromolecular composition analysis. Samples for phycocyanin quantification were collected daily. Seasonal cultivation conditions representative of Thuwal, Saudi Arabia (22.3046 N, 39.1022 E), were simulated in Algem photobioreactors using environmental profiles previously reported (Freitas et al., 2023). Simulations corresponded to February, May, August, and November conditions, emulating Winter, Spring, Summer, and Autumn, respectively. Biomass composition was assessed using samples collected between d4-7, and phycocyanin content was monitored daily.

### 2.4 Biomass macromolecular composition

Biomass sampels were frozen at -80 °C, freeze-dried, and stored at -20 °C until further analysis. For carbohydrate and protein analysis, intracellular material was released from 5 mg of dried biomass suspended in 10 mL distilled water using an ultrasonic probe (FB705, Fisher Scientific, Pittsburgh, PA, USA) for 10 min.

Total carbohydrates were quantified using the phenol-sulfuric acid method with glucose as the standard (DuBois et al., 1956). Briefly, 1 mL of biomass extract was mixed with 1 mL of 5 % phenol solution, followed by addition of 5 mL concentrated H_2_SO_4_ Samples were incubated for 10 min, cooled in a cold-water bath for 20 min, and absorbance was measured at 488 nm. Glucose was used as standard.

Total protein content was determined using the Lowry method (Lowry et al., 1951). Biomass extract was mixed with 1 M NaOH and incubated at 100 °C for 10 min, cooled for 10 min, and reacted with alkaline copper reagent and Folin solution. Samples were incubated for 30 min in the dark and absorbance was measured at 750 nm. Bovine serum albumin was used as standard.

Total lipid content was determined from 10 mg of dried biomass using the colorimetric method described by Marsh and Weinstein (Marsh & Weinstein, 1966). Biomass disruption was performed using a FastPrep-24 5G (MP Biomedicals) with Lysing Matrix D tubes for three cycles of 60 s, using chloroform:methanol (1:2, v/v). The extract was transferred to glass tubes, heated at 60 °C for 5 min, and repeatedly extracted with chloroform:methanol (1:2, v/v) until the pellet and supernatant were colorless. Phase separation was performed by adding chloroform and Milli-Q water, and the lower organic phase was collected and dried at 80 °C. Dried lipid extracts were resuspended in chloroform, aliquoted, dried again, reacted with H_2_SO_4_ at 200 °C for 15 min, diluted with Milli-Q water, and absorbance was measured at 375 nm. Tripalmitin was used as standard (Holland & Gabbott, 1971).

### 2.5 Pigment extraction

Chlorophyll a concentrations and total carotenoids were calculated according to Arnon (Arnon, 1949) and Lichtenthaler and Buschmann (Lichtenthaler and Buschmann, 2005), respectively, from 10 mg of dried biomass using 1 mL of 80% acetone and the same bead-beating procedure drescribed above. Extracts were diluted to a final volume of 10 mL. Absorbance readings were recorded at 470 nm, 645 nm, and 663 nm.

Pigment extractions were performed under dim light conditions. Freeze-dried biomass (10 mg) was mixed with 200 mg of 1 mm glass beads and extracted using either 1.5 mL of 0.1 M phosphate buffer (pH 7.0) for C-PC extraction or acetone:water (9:1 v/v - for carotenoid and chlorophyll extraction) precooled to 4°C and ran for 60 s at 6 m s^-^1 for 3 cycles with 60 s in bead mill as above with intermediate cooling on ice. Extracts were centrifuged at 12,000 × g for 3 min and supernatants collected. Pellets were re-extracted until colorless and solvents pooled before analysis, acetone extracts were evaporated under nitrogen.

For carotenoid saponification, pigment extracts were dried under N_2_, resuspended in hexane, and incubated with 5 % (w/v) methanolic KOH for 2 h at room temperature under dim light. Saponification was stopped by adding 100 µL of 10 % (w/v) NaCl and 200 µL of deionized water. The unsaponified carotenoid fraction was recovered by four sequential extractions with hexane:MTBE (1:1 v/v; 300 µL per extraction), with phase separation by centrifugation at 12,000 × g for 1 min after each extraction. The organic fractions were pooled, dried under nitrogen, resuspended in 1 mL acetonitrile, and filtered through 0.45 µm nylon filters before analysis by TLC, UV-Vis spectrophotometry, and HPLC. Non-saponified extracts were processed in parallel by drying, resuspension in acetonitrile, and filtration prior to analysis.

Thin-layer chromatography (TLC) was performed using silica gel 60 plates (Supelco) with hexane:acetone (7:3, v/v) as the mobile phase. Carotenoid standards were obtained from CaroteNature GmbH (Munsingen, Switzerland). The standard panel included ß-carotene (No. 0003), echinenone (No. 0283), canthaxanthin (No. 0380), ß-cryptoxanthin (No. 0055), adonirubin (No. 0391), astaxanthin (No. 0403), adonixanthin (No. 0328), isozeaxanthin (No. 0129), violaxanthin (No. 0259), and lutein (No. 0133; containing approximately 5% zeaxanthin). Unless otherwise indicated, standards were supplied as crystalline compounds with HPLC purity ≥95%.

### 2.6 High-Performance Liquid Chromatography - Photodiode Array detector carotenoid analysis

Carotenoids were separated and quanitified using a Thermo Scientific Vanquish Ultra High-Performance Liquid Chromatography (UHPLC) system coupled to a photodiode array (PDA) detector. Pigments were separated on an Acclaim™ 120 C18 column (5 μm, 120 Å, 4.6 × 100 mm) maintained at 25 °C. The mobile phase consisted of ethyl acetate and acetonitrile:water:triethylamine (9:1:0.01, v/v/v). Carotenoids were separated using a 28 min gradient of ethyl acetate (0–100%) at 1.0 mL min^-1^.

Chromatograms were monitored at 430, 450, 470, and 480 nm, and absorbance spectra were acquired from 280 to 700 nm with the PDA detector. Data acquisition, peak integration, and spectral analyses were carried out using Chromeleon v. 7.3.1. Peak identities were assigned by comparison with authentic standards and quantified using external calibration curves. Peak purity was assessed by comparing PDA spectra across each chromatographic peak using Chromeleon v. 7.3.1. Peak assignments were considered valid when retention times and absorbance spectra matched those of authentic standards.

### 2.7 Phycocyanin extraction and stability

Freeze-dried biomass (∼50 mg) was resuspended in 1 mL PBS buffer (pH 7.4) precooled to 4 °C and incubated for 1 h with continuous shaking. Samples were centrifuged at 10,000 × g for 10 min and the supernatants were collected for spectrophotometric analysis. C-phycocyanin (C-PC) and allophycocyanin (A-PC) concentrations were determined spectrophotometrically using absorbance measurements at 615, 652, and 720 nm with a Genesy™ 50 UV-Vis spectrophotometer (Thermo Scientific, UK). Concentrations were calculated according to Equations 1 and 2.

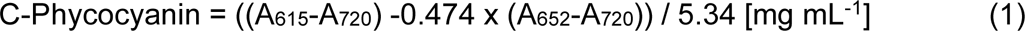

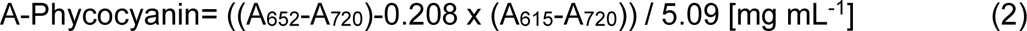

The thermostability of extracts obtained from *Limnospira platensis* and the *Halospirulina* isolate were evaluated after normalization to 1 mg mL^-1^ C-phycocyanin. Aliquots (50 µL) were exposed to the indicated temperature treatments using a T100 Thermal Cycler (Bio-Rad, USA). Following incubation, samples were cooled on ice prior to analysis. For image acquisition, four 50 µL aliquots from each temperature treatment were pooled into a single tube to improve visualization of color changes.

## 3 Results

### 3.1 Isolation and identification of *Halospirulina saudiensis* sp. nov

A filamentous cyanobacterial strain was isolated from samples collected at a water-clay microhabitat (pH 6.8) in the Empty Quarter desert, Saudi Arabia as described above (Figure 1A and B). This geography is a hyper-arid region characterized by intense irradiance, elevated and extreme diurnal fluctuation in temperatures, as well as severe water limitation (Blanco-Sacristán et al., 2025; Edgell, 2006). Enrichment cultures were maintained at 42 ± 3 °C using Red Sea water-based medium for 14-20 d. Cultures were periodically examined by light microscopy, and serial dilution was performed until a cyanobacterial monoculture was established. Microscopy revealed unbranched, helically coiled trichomes characteristic of members of the *Spirulinaceae* (Figure 1D) which lead us to further investigate its identity (Komárek et al., 2014).

**Figure 1.**
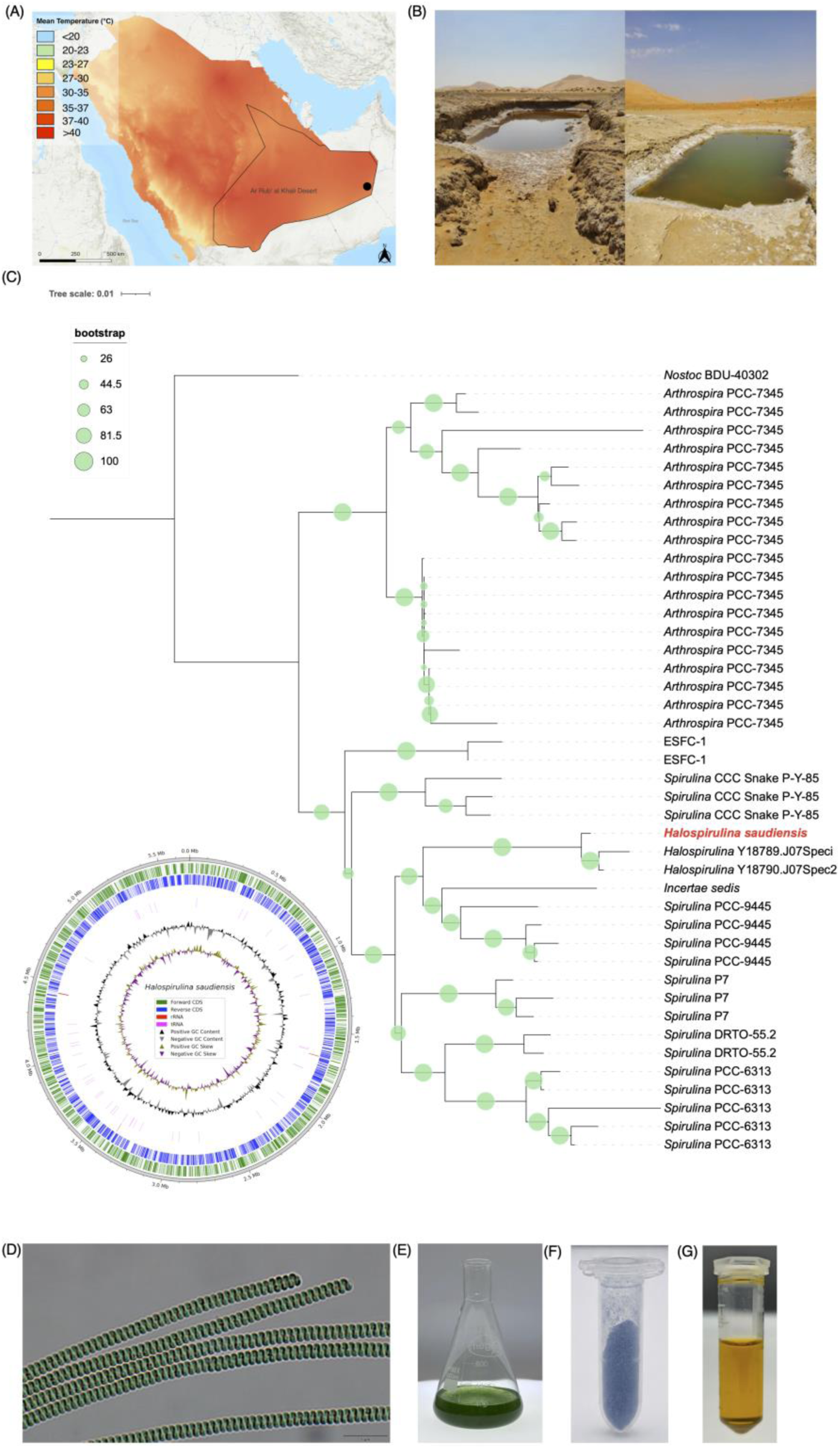
Isolation and characterization of *Halospirulina saudiensis* sp. nov. (A) Mean annual temperature profile of the Arabian Peninsula indicating the Empty Quarter (Rub’ al Khali), Saudi Arabia, where sampling was conducted (black dot). (B) Sampling site showing the shallow saline water microhabitat from which the strain was isolated (Hoehndorf, 2026). Maximum likelihood phylogenetic tree based on 16S rRNA sequences showing the placement of *H. saudiensis* within *Spirulinaceae*. Circle size indicates bootstrap support values. The inset shows the circularized genome assembly of *H. saudiensis*, including coding sequences and GC content. *Nostoc* BDU-40302 was used as the outgroup. (D) Light microscopy image showing the helically coiled trichomes of *H. saudiensis*. (E) Biomass cultivated in BG-11 prepared with ultrafiltered Red Sea seawater. (F) Extracted total soluble protein containing C-phycocyanin and freeze dried to powder. (G) Carotenoid extract (acetone) obtained from *H. saudiensis* biomass.

Taxonomic identification was performed using a polyphasic approach integrating morphology, 16S rRNA phylogeny, whole-genome phylogenomics, and genome similarity analysis (Komárek et al., 2014). Taxonomic classification followed current systematic revisions of *Spirulinales* and the Madrid Code (Turland, 2025). Whole-genome sequencing generated a circularized ∼5.7 Mb genome assembly (NCBI Accession No. JBYDFK000000000, and Supplementary material 2). Analysis of 16S rRNA phylogeny placed this isolate close to available *Halospirulina* sequences (Figure 1C) and a phylogenomic tree separated it from available *Limnospira* and *Arthrospira* genomes (Supplementary Figure S1). This fully sequenced genome represents to the best of our knowledge, the first available for a *Halospirulina*. Genome similarity analysis showed a maximum ANI of 83 % relative to the closest available reference genome (Supplementary Figure S2), indicating substantial genomic divergence from *Spirulina* sp. CS-785/01. Based on its phylogenetic placement, genome divergence, morphology, and geographic origin, we propose the name *Halospirulina saudiensis* sp. nov.

The isolate, *H. saudiensis*, morphologically exhibits tricome spirals similar to *Limnospira* (Figure 1D), but was enriched using saline cultivation conditions (Figure 1E), while surprisingly exhibiting a carotenoid profile containing C-PC pigment (Figure 1F) and ketocarotenoids (Figure 1G). These traits warranted further exploration as to whether it may be interesting as a species for biomass and bioproduct generation in saline cultivation concepts.

### 3.2 Physiological response to pH and temperature

Following isolation to monoculture, cultivation experiments were performed to evaluate the physiological response of *H. saudiensis* under controlled conditions. Cultures were initially grown in photobioreactors at 40 °C under continuous illumination (325 µmol photons m^-2^ s^-1^), corresponding to the conditions used during enrichment and isolation. Preliminary experiments evaluated growth at pH 8, 9, and 10, as well as under continuous CO_2_ supplementation (Supplementary Figure S3). Maximum biomass was observed at pH 8 (5.09 g L^-1^ after 7 d), while pH 9 cultures struggled to grow (2.18 g L^-1^). At pH 10, cultures lost viability within 3 d. Continuous CO_2_ supplementation resulted in lower maximum biomass (<1.5 g L^-1^), followed by a decline after d5. Based on these observations, subsequent experiments were performed between pH 7-9 at either 35 or 40 °C to determine the optimal growth conditions.

Biomass accumulation was strongly influenced by temperature and pH. At 35 °C, cultures grown at pH 7 and 8 reached 9-10 g L^-1^ by d4, whereas pH 9 exhibited lower biomass overall and declined faster with cultures showing visible clumping (Figure 2B). At 40 °C, biomass was slightly lower overall, with the highest values observed at pH 8 (8.38 ± 0.77 g L^-1^) (Figure 2B). Cultures at pH 7 reached maximum biomass between d5-6 (9.83 g L^-1^) using 35 °C, pH 7, and 325 µmol m^-2^ s^-1^ (Figure 2B). This biomass yield exceeds values reported for some *Arthrospira/Spirulina* cultivation systems under controlled photobioreactor conditions, which typically range between 1.5 and 3.0 g L^-1^ (Barahoei et al., 2026; Ma et al., 2025; Maneechote et al., 2025).

**Figure 2.**
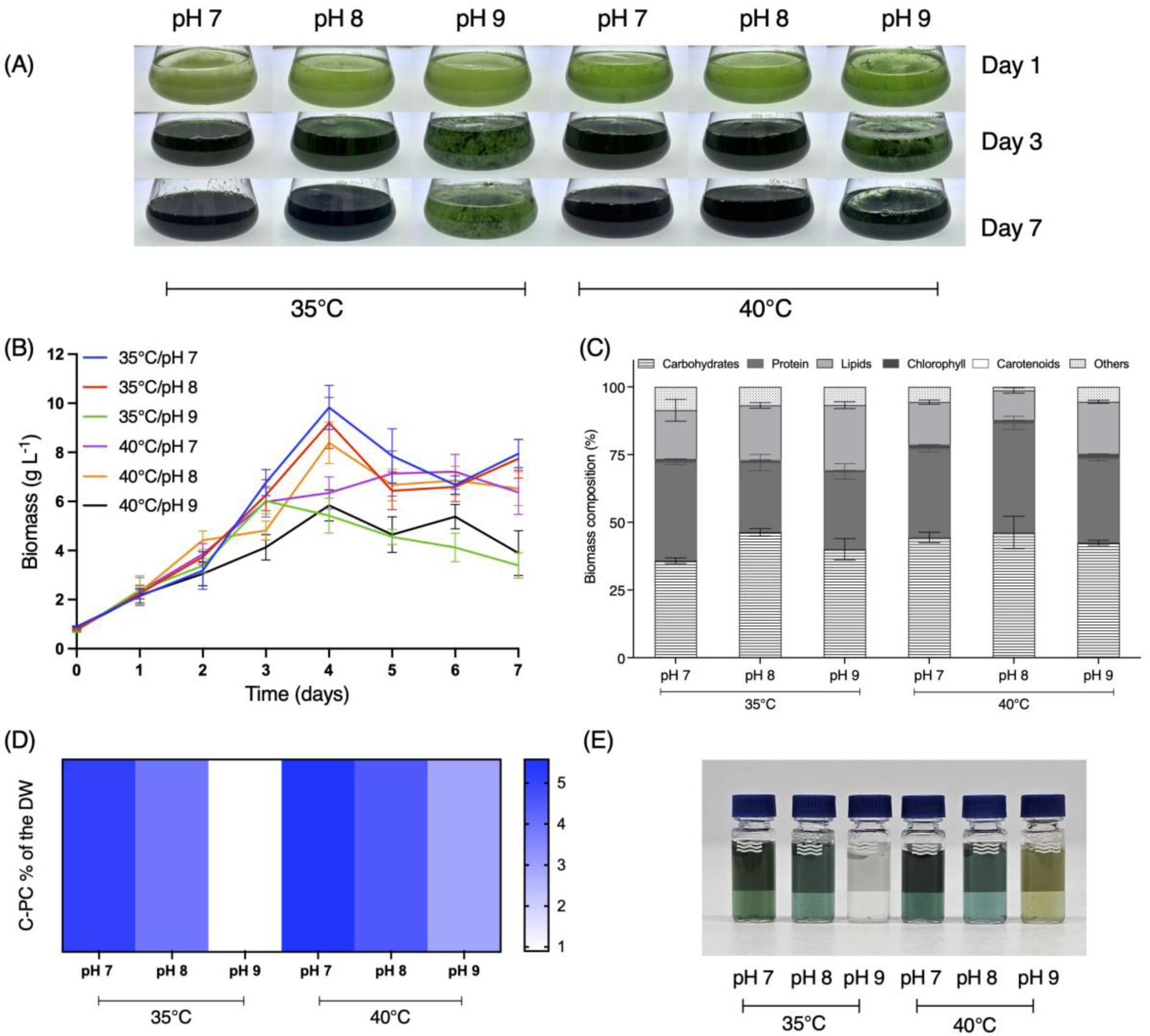
Effect of pH and temperature on growth performance, biomass composition, and C-PC accumulation of *H. saudiensis*. (A) Culture appearance during cultivation at pH 7, 8, and 9 under 35 and 40 °C during cultivation (d1, 3, and 7). (B) Biomass accumulation (g L^-1^) during 7d of cultivation under different pH and temperature conditions. Values represent mean ± SD of biological duplicates (n = 2), with biomass determined from 3 technical measurements per replicate. (C) Biomass composition (% dry weight) determined at the end of cultivation. Values represent mean ± SD of biological duplicates (n = 2). (D) C-PC content expressed as percentage of dry biomass under the evaluated cultivation conditions. Values represent mean ± SD (n = 3) (E) C-PC extracts obtained from biomass cultivated under the indicated conditions.

Under the highest biomass-producing condition (35 °C, pH 7), biomass composition was characterized by balanced carbohydrate and protein fractions (35.8 and 36.5 %, respectively), moderate lipid content (17.9 %), and comparatively low chlorophyll and carotenoid fractions (<1 % DW). In contrast, cultures grown under pH 8 conditions showed higher carbohydrate fractions (46.3%) and lower protein contents (25.7%) despite maintaining high biomass accumulation (Figure 2C). C-phycocyanin (C-PC) content was also influenced by pH and temperature (Figure 2D). C-PC accumulation was highest at pH 7 under both temperature conditions (5.1-5.6 %) and remained relatively high at pH 8 (3.9-4.5 %), whereas pH 9 consistently resulted in lower C-PC content.

### 3.3 Effect of irradiance on growth and C-PC production

Based on the pH and temperature screening, pH 7, 35 °C were selected as base parameters and additional irradiance levels were applied to evaluate the effect of light intensity on growth and biomass composition. Biomass accumulation varied with irradiance and cultivation time (Figure 3A). Maximal biomass was observed under 325 µmol photons m^-2^ s^-1^ on d4, whereas cultures grown under 200 and 500 µmol photons m^-2^ s^-1^ reached maximal biomass at later cultivation stages between d6-7 (Figure 3A). Following peak biomass accumulation, stationary phase cells progressively accumulated carbohydrates, an effect that was most pronounced under high irradiance. Under 500 µmol photons m^-2^ s^-1^, carbohydrate content reached 56.3% at d7, while protein content decreased to ∼20%. In contrast, cultures grown under 200 µmol photons m^-2^ s^-1^ maintained protein contents above 45% throughout the later cultivation stages. Lipid content varied across irradiance conditions without a consistent temporal trend and ranged between 11.4 - 25.7% of biomass (Figure 3B). C-PC content varied with both irradiance and cultivation time (Figure 3C). At 200 µmol photons m^-2^ s^-1^, C-PC increased toward the end of cultivation, rising form 12.1 to 65.9 mg g^-1^ DW between d1 and d7. At 325 µmol photons m^-2^ s^-1^, C-PC increased rapidly after d2, reaching a maximum of 62.5 mg g^-1^ DW at d6 before declining to 44.4 mg g^-1^ DW at d7. In contrast, cultures grown at 500 µmol photons m^-2^ s^-1^ showed variable C-PC levels, with maximum values (67.3 mg g^-1^ DW) observed at intermediate cultivation stages.

**Figure 3.**
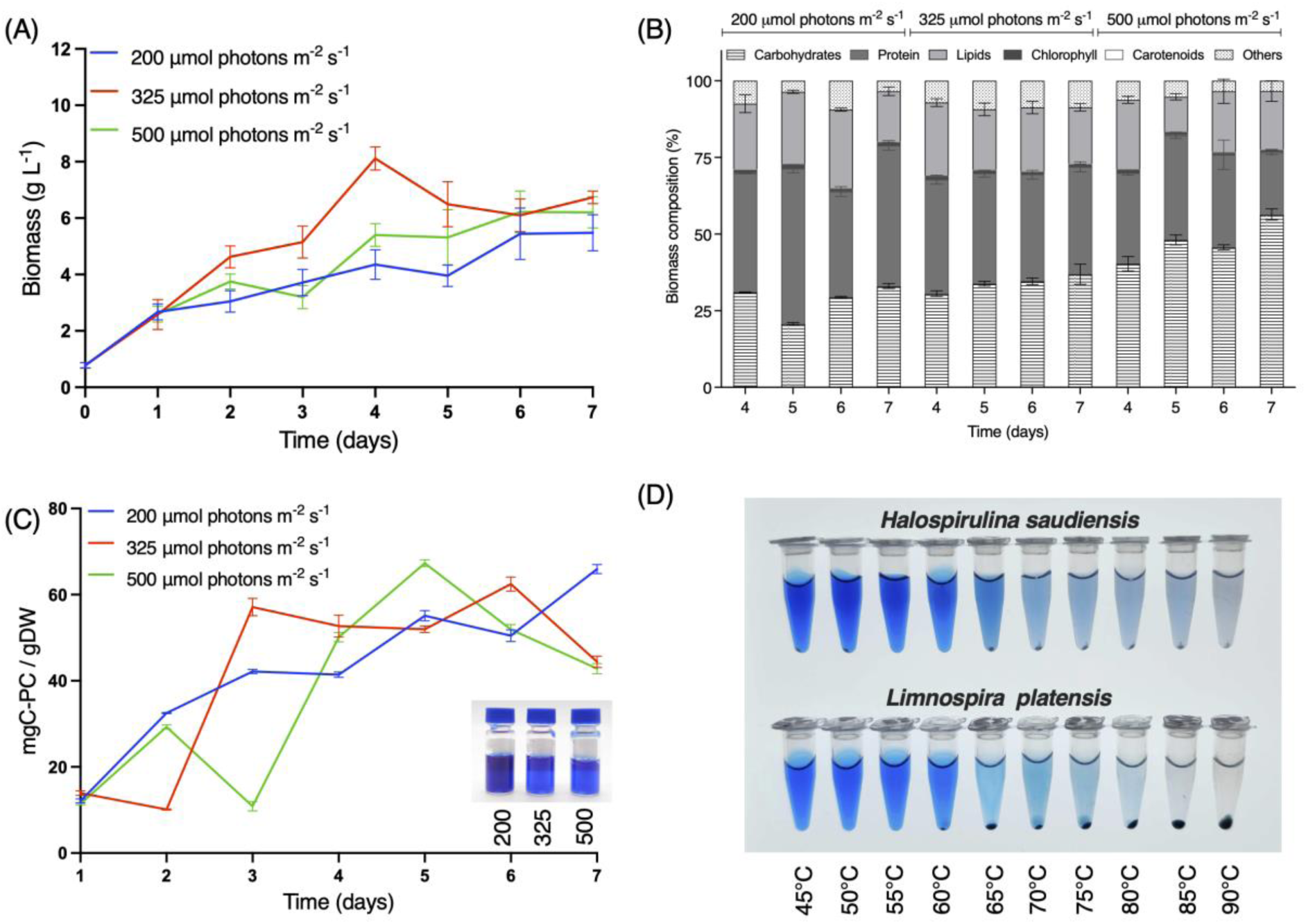
Effect of irradiance on biomass accumulation, biomass composition, C-PC accumulation, and phycocyanin stability of *H. saudiensis*. (A) Biomass accumulation (g L^-1^) during 7d of cultivation under 200, 325, and 500 µmol photons m^-2^ s^-1^. Values represent mean ± SD of biological duplicates (n = 2), with biomass determined from 3 technical measurements per replicate. (B) Biomass composition (% dry weight) determined from biomass collected between d4-7 under different irradiance conditions. Values represent mean ± SD of biological duplicates (n = 2). (C) C-PC content expressed as mg C-PC g^-1^DW during cultivation under different irradiance conditions. Values represent mean ± SD (n = 3). C-PC extracts are shown in the inset. (D) Thermal stability of extracted C-PC from *H. saudiensis* and *L. platensis* after incubation between 45 and 90 °C.

### 3.4 Phycocyanin stability

Total soluble proteins were extracted from biomass of *H. saudiensis* and *L. platensis*. Clear blue supernatants were obtained for both strains, with no detectable residual pigmentation after 3 extractions. Following normalization, thermal stability assays show progressive loss of blue color in both extracts, however, *H. saudiensis* retained visibly stronger coloration at elevated temperatures compared to *L. platensis* (Figure 3D). Differences became evident above 65 °C, where *L. platensis* showed color loss, while *H. saudiensis* retained blue coloration, although reduced in vibrancy at higher temperatures. Enhanced thermal stability of phycocyanin has previously been reported in extremophilic microalgae and cyanobacteria adapted to elevated temperature conditions (Masson et al., 2025; Wang et al., 2025).

### 3.5 Ketocarotenoid accumulation in *H. saudiensis*

The coloration of *H. saudiensis* primary acetone extract differed markedly from that of *L. platensis* (Figure 1G). Therefore, the carotenoid profile of *H. saudiensis* was analyzed using biomass cultivated under its optimally identified conditions (pH 7, 35 °C, above) and compared to *L*. platensis (Figure 4A). The HPLC chromatogram of *L. platensis* was dominated by ß-carotene, zeaxanthin, and two major peaks tentatively assigned as hydroxyechinenone-related carotenoids, together with early-eluting xanthophyll-like compounds. In contrast, *H. saudiensis* lacked the prominent zeaxanthin associated peaks observed in *L. platensis* and instead showed increased abundance of secondary ketocarotenoids. Major peaks included canthaxanthin, a putative cis-canthaxanthin-like carotenoid (λ_max_366 and 468 nm), a putative myxoxanthophyll-like carotenoid (λ_max_296, 476, and 507 nm), and a carotenoid eluting in the echinenone region. Differences in carotenoid composition were also visible in the saponified extracts, where *H. saudiensis* displayed a more orange coloration than *L. platensis* (Figure 4A). A major carotenoid peak detected in both strains eluted at the same retention time as an authentic echinenone standard and co-migrated with the standard by TLC (Supplementary Figure S4). However, its PDA absorption spectrum differed from that of authentic echinenone, preventing definitive assignment of the compound as pure echinenone. This pigment was, therefore, conservatively annotated as an echinenone-like carotenoid.

**Figure 4.**
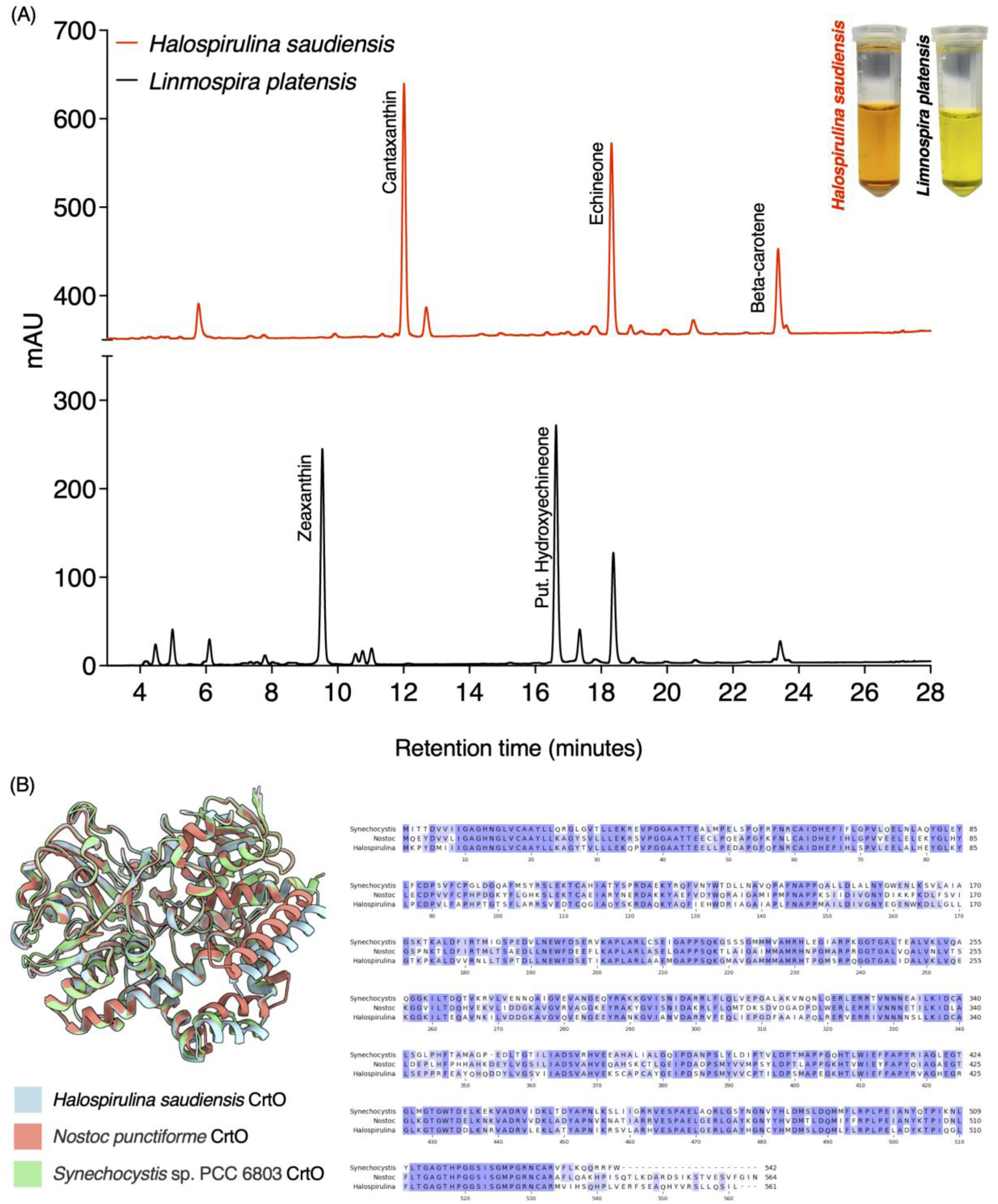
Ketocarotenoid accumulation and CrtO characterization in *H. saudiensis*. (A) HPLC chromatograms of carotenoid extracts from *H. saudiensis* and *L. platensis* cultivated under their respective cultivation conditions. Major peaks are annotated according to putative or confirmed pigment identity. Insets show pigment extracts after saponification. (B) Structural comparison of predicted CrtO proteins from *H. saudiensis*, *N. punctiforme*, and *Synechocystis* sp. PCC 6803. Protein structural overlays and multiple sequence alignment are shown.

*In silico* modelling showed high structural similarity between the predicted *H. saudiensis* CrtO protein and homologs from *Nostoc punctiforme* and *Synechocystis* sp., with root mean square deviation (RMSD) values of 0.420 Å and 0.528 Å, respectively. These homologs were selected based on their established roles in cyanobacterial ketocarotenoid biosynthesis (Fernandez-Gonzalez et al., 1997). Structural overlays and sequence alignments are shown in Figure 4B and support the overall conservation of the CrtO homologs, although functional validation of its role in keto carotenoid biosynthesis is still required.

### 3.6 Growth performance under simulated seasonal environments

Seasonal cultivation conditions representative of February, May, August, and November in the mid Red Sea coast of Saudi Arabia, were simulated in photobioreactors using environmental temperature and irradiance profiles (Figure 5A) following the approach previously described by de Freitas et al. (2023). Biomass accumulation varied across seasonal profiles, with the highest values observed under August (4.7 g L^-1^, d7) and May (4.4 g L^-1^, d7) conditions, whereas February consistently resulted in lower biomass throughout cultivation (Figure 5B). Biomass composition also varied across seasonal conditions (Figure 5C). Protein content was highest in May (44.2%) and August (40.2%), whereas February and November showed lower values (37.5 and 34.7, respectively). Lipid content peaked in November (28.95%) and was lowest in May (19.1%). In contrast, carbohydrate levels remained relatively stable across all conditions (28.4-32.2 %). C-PC accumulation was strongly reduced under February conditions, remaining below 10 mg gDW^-1^ throughout cultivation (Figure 5D). In contrast, May, August, and November reached similar maximum C-PC levels ranging from 54.5-61.1 mg gDW^-1^ during the later cultivation stages.

**Figure 5.**
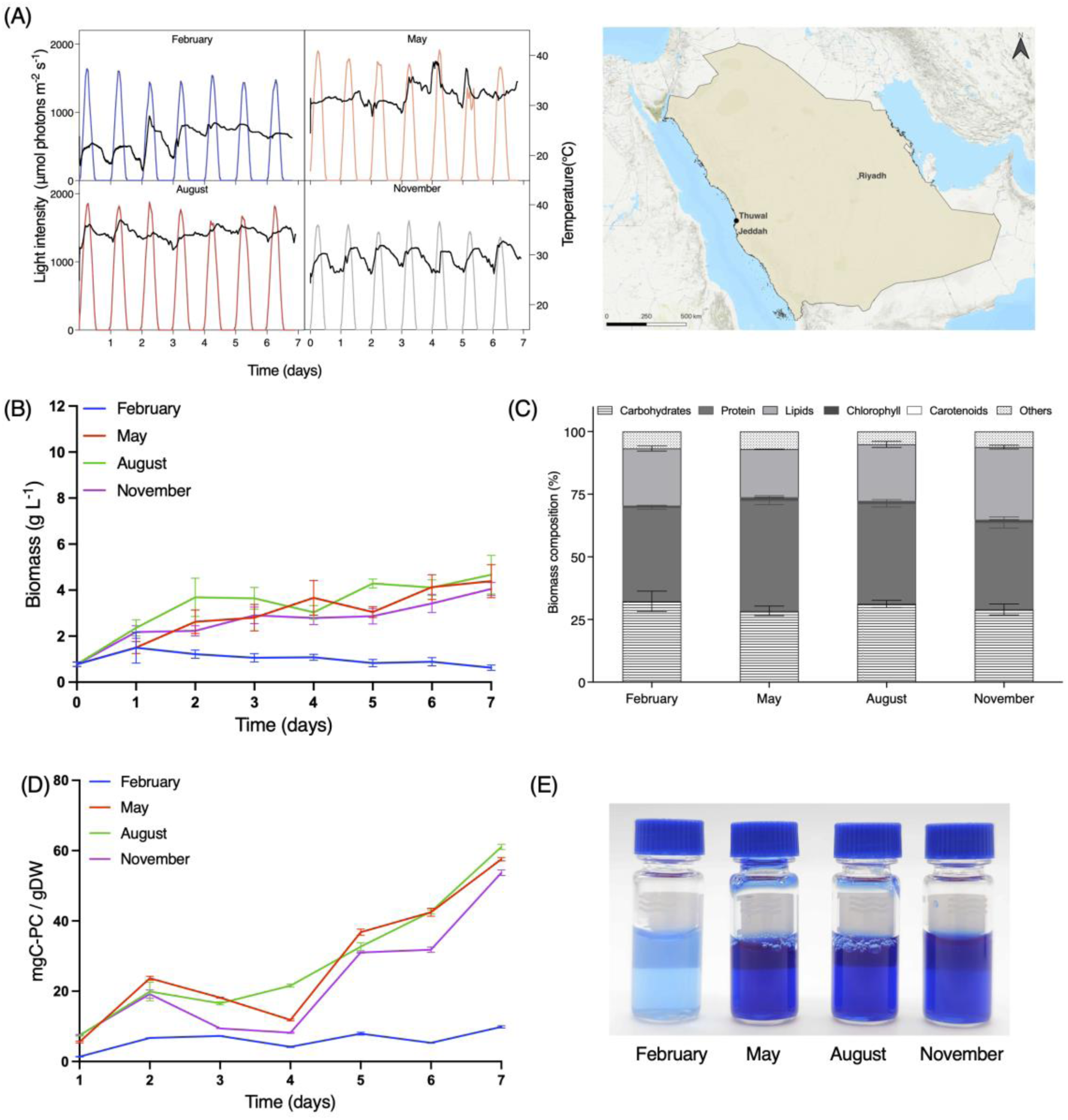
Effect of simulated seasonal conditions on biomass accumulation, biomass composition, and C-PC accumulation of *H. saudiensis*. (A) Simulated irradiance and temperature profiles representative of February, May, August, and November conditions in Thuwal, Saudi Arabia. The map indicates the location used as reference for environmental simulations. (B) Biomass accumulation (g L^-1^) during 7d of cultivation under simulated seasonal conditions. Values represent mean ± SD of biological duplicates (n = 2), with biomass determined from 3 technical measurements per replicate. (C) Biomass composition (% dry weight) determined at the end of cultivation under each seasonal condition. Values represent mean ± SD of biological duplicates (n = 2). (D) C-PC content expressed as mg C-PC g^-1^DW during cultivation under simulated seasonal conditions. Values represent mean ± SD (n = 3). (E) C-PC extracts from biomass cultivated under each seasonal condition.

## 4 Discussion

The taxonomy of filamentous cyanobacteria historically classified within *Spirulina*-related lineages remains complex, particularly due to the limited availability of genome-resolved representatives for several environmental *Spirulinaceae* groups (Komárek et al., 2014; Nowicka-Krawczyk et al., 2019; Nübel et al., 2000). Here, 16S rRNA phylogeny associated our isolate with available *Halospirulina* sequences and phylogenomic analysis clearly separated the strain from currently available *Limnospira* and *Arthrospira* genomes. Although ANI comparisons, the gold standard for claims of newly identified species, are limited here by the absence of additional Halospirulina genomes, the morphology, phylogenetic placement, genome divergence, and ecological origin of *H. saudiensis* support its recognition as a distinct *Halospirulina* lineage (Chun et al., 2018; Komárek et al., 2014). The circularized genome generated in this study represents the first genome available for a *Halospirulina* type strain (NCBI Accession No. JBYDFK000000000, and Supplementary material 2).

In our lab-scale cultivations, *H. saudiensis* maintained high biomass accumulation in Red Sea seawater-based media, which has higher salinity that global average, and under environmentally simulated temperature and irradiance regimes. Although growth was maintained at pH 8 and tolerance to pH 9 was observed, the strain did not exhibit the strong alkalitolerance commonly associated with industrially cultivated *Limnospira* systems (Li et al., 2024; Vasquez Guevara et al., 2025). The strain was isolated from wet clay with a pH of 6.8, so was not naturally in an alkali environment. Genome analysis identified homologs associated with compatible solute biosynthesis, including glucosylglycerol-phosphate synthase (*ggpS)* and glucosylglycerol-phosphate phosphatase (*stpA)*, together with additional putative homologs of trehalose-6-phosphate synthase *(otsA)* and sucrose-phosphate synthase (*sps)*, together with conserved ion homeostasis systems including the low-affinity Na⁺/H⁺ antiporter (*nhaS1*) and components of the multiple resistance and pH adaptation (*mrp*) Na^+^/H^+^ antiporter complex. These systems are commonly associated with osmotic stress acclimation and sodium homeostasis in cyanobacteria inhabiting saline environments (Hagemann and Marin, 1999; Hamada et al., 2001; Kirsch et al., 2019; Marin et al., 1998; Waditee-Sirisattha and Kageyama, 2023)

The combined physiological and genomic observations suggest that salinity adaptation may play a more important role in the physiology of *H. saudiensis* than extreme alkaliphily. These traits are consistent with the isolation of *H. saudiensis* from a hypersaline desert-associated microhabitat exposed to strong irradiance and elevated temperature fluctuations (Hoehndorf, 2026). Irradiance strongly influenced biomass composition and C-PC accumulation. Cultures grown under elevated irradiance accumulated larger carbohydrate fractions during later cultivation stages, whereas lower irradiance maintained higher protein contents and sustained C-PC accumulation. Similar shifts in biomass composition under different cultivation conditions have previously been reported in microalgae and cyanobacteria (Arenas Colarte et al., 2025; Oslan et al., 2025). Overall, C-PC accumulation was not directly coupled to biomass formation, and reduced C-PC levels under higher irradiance were consistent with previously reported photoprotective responses in cyanobacteria exposed to excess light (Sanfilippo et al., 2019; Tamary et al., 2012).

In addition to differences in phycobiliprotein accumulation, *H*. *saudiensis* displayed a carotenoid profile enriched in secondary ketocarotenoids relative to *L. platensis*. Genome analysis identified a *crtO* homolog related to those reported in *Nostoc punctiforme* and *Synechocystis* sp. PCC 6803, both of which contain carotene ketolases associated with echinenone and canthaxanthin biosynthesis (Breitenbach et al., 2013; Fernández-González et al., 1997; Kłodawska et al., 2019). Ketocarotenoids such as canthaxanthin and echinenone are commonly associated with antioxidant and photoprotective functions in cyanobacteria (Kłodawska et al., 2019; Llewellyn et al., 2020; Pagels et al., 2021). The increased abundance of these pigments and reduced zeaxanthin distinguishes the carotenoid profile of *H. saudiensis* from that of *L. platensis*. This profile is consistent with the accumulation of photoprotective ketocarotenoids previously associated with high-light acclimation in cyanobacteria (Breitenbach et al., 2013; Llewellyn et al., 2020).

Beyond their proposed roles in photoprotective acclimation, ketocarotenoids such as canthaxanthin are industrially relevant pigments used in aquaculture, feed, cosmetic, and nutraceutical applications due to their antioxidant and coloration properties (Chakdar and Pabbi, 2017; Rebelo et al., 2020; Udayan et al., 2017). Although the commercial supply of canthaxanthin currently relies predominantly on chemical synthesis, growing consumer awareness regarding health and environmental safety is driving a strong industrial demand for natural alternatives (Oslan et al., 2025; Rebelo et al., 2020). In this context, the capacity of the isolate to naturally synthesize these highly valued compounds provides a safer and more sustainable substitute (Mahanandia et al., 2025). The combined accumulation of ketocarotenoids together with the stability of extracted phycocyanin suggests that *H. saudiensis* may represent an interesting candidate for pigment-oriented cultivation and bioprocessing under saline and elevated temperature cultivation conditions (Bounnit et al., 2022; Mahanandia et al., 2025).

The extracted phycocyanin fraction from *H. saudiensis* showed lower loss of visible blue coloration at temperatures above 65 °C compared with *L. platensis*, although color degradation was observed in both strains under these conditions. Phycocyanin from commercial *Spirulina/Limnospira* strains is generally considered thermolabile (Antecka et al., 2022; Chaiklahan et al., 2012). Pasteurization is possible at 65 °C, which means C-PC from *H. saudiensis* would not require stabilization prior to pasteurization and could be used in a more natural state. Although the molecular basis underlying this phenotype remains unresolved, increased thermal stability of the extracted pigment fraction may be associated with its tolerance to elevated environmental temperatures (Liang et al., 2018; Palanivel et al., 2026).

The western coast of Saudi Arabia has recently been identified as a favorable region for large-scale microalgae cultivation due to the combination of high solar irradiance, extensive non-arable land, direct seawater access, and proximity to industrial CO_2_ sources (Mhedhbi et al., 2026). Within this favorable cultivation context, *H. saudiensis* exhibited seasonal variation in biomass productivity and biomass composition that was closely associated with temperature and irradiance. Here, seasonal cultivation simulations of *H. saudiensis* were performed in BG-11 prepared with undiluted Red Sea seawater, indicating that the high biomass production was maintained under the elevated salinity marine medium conditions relevant to the western Saudi Arabian coast (Krokos et al., 2024).

February conditions consistently resulted in reduced biomass accumulation and C-PC production, whereas May and August supported higher protein accumulation and biomass productivity. Outdoor cultivation of *Limnospira maxima* in Thuwal has also shown that warmer periods support higher biomass productivity and protein-rich biomass under Red Sea coastal conditions (up to 0.28 g L^-1^ d^-1^)(González-Portela et al., 2024). Although direct comparison between cultivation systems should be interpreted cautiously, our simulation of seasonal conditions representative of Thuwal in photobioreactors supported higher biomass accumulation (up to 1.46 g L^-1^ d^-1^) than previously reported for outdoor raceway cultivation. This result suggests that *H*. *saudiensis* maintains high productivity under environmentally-relevant outdoor cultivation conditions. Similar seasonal effects on biomass productivity, protein accumulation, and C-PC content have also been reported in *Limnospira* cultivation systems, where Spring and Summer conditions supported higher protein and phycocyanin levels (Lopes et al., 2026).

The Arabian Peninsula, as well as many other coastal desert geographies, exhibit routine severe climatic events, such as heat waves and dust storms which carry with them uncharacterized microbial contaminants (Blanco-Sacristán et al., 2025). Additionally, local fauna, especially birds, may frequent open pond cultivations, these factors may introduce contamination risks in large, open, cultivation systems, especially if long-standing cultures are maintained (Guidi et al., 2021; Molina-Grima et al., 2022). These risks are minimized in classic *Limnospira* culture, due to the incredibly alkaline pH extremes of this species (Nowicka-Krawczyk et al., 2019; Vasquez Guevara et al., 2025). Even greater biomass was achieved here with 24 h illumination (9.83 g L^-1^). We found that despite its interesting bioproducts and saline cultivation preference, *H. saudiensis* is not an alkaliphile, but rather must be cultivated at neutral pH. These parameters suggest it would be favorable to cultivate it in continuous, contained photobioreactor systems to minimize contamination. As sun-light is abundant in regions like the Middle East, reactors could couple daytime sun-light driven illumination with augmentation through artificial illumination during the night to reduce operating costs and increase biomass output. Tubular photobioreactors, or illumination supported wall panels, would be attractive options. The exhibited thermotolerance of *H. saudiensis* due to its origin from the Empty Quarter, likely will reduce cooling needs in scaled cultures.

## 5 Conclusions

This study presents the first genome-resolved characterization of a *Halospirulina* type strain. Our isolate, comes from the Empty Quarter desert of Saudi Arabia, a region known for high irradience, daily, diurnal and seasonal temperature fluctuations, and low water availability. *H. saudiensis* maintained growth under saline cultivation conditions prepared using Red Sea seawater across neutral-slightly alkali pH, temperature, irradiance, as well as simulated seasonal cultivation profiles representative of the central western coast of Saudi Arabia. Biomass concentrations reached 9.8 g L⁻¹ within 4 d in continuous controlled cultivation, while maximum C-phycocyanin concentrations reached 67.3 ± 0.8 mg gDW⁻¹ under higher irradiance conditions. Biomass production was also maintained under simulated May and August outdoor environmental conditions, reaching up to 4.7 g L^-1^. In addition to biomass production, *H. saudiensis* displayed an unusual carotenoid profile containing secondary ketocarotenoids with the highest abundance in canthaxanthin, currently an antioxidant pigmentation additive used in chicken and fish feeds. The unique carotenoid profile of *H. saudiensis*, coupled to production of C-PC with improved thermostability also means the biomass can be used in biorefinery concepts, yielding blue soluble C-PC extracts and oil-soluble ketocarotenoids. The combination of rapid biomass accumulation under saline cultivation conditions with such biomass fractions suggests that *H. saudiensis* may represent a promising candidate for large scale cultivation in high temperature coastal environments as found in the Middle East and North Africa. But its neutrophilic nature likely requires contained cultivation systems in order to ensure stable, contaminant free production.

## Supporting information

Supplementary material 1 - Protein sequences

Supplementary material 2 - Genome assembly

Supplementary material 5 - Biomass composition

Supplementary material 6 - C-phycocyanin

Supplementary material 7 - Chromatogram HPLC Halospirulina

## 6 Taxonomic Note: *Halospirulina saudiensis* Bastos de Freitas & Lauersen, sp. nov

### Etymology

sau.di.en’sis. L. fem. adj. saudiensis, referring to the Kingdom of Saudi Arabia, where the strain was isolated.

### Diagnosis

Distinguished from other Spirulinaceae cyanobacteria by its isolation from a hyper-arid desert water-clay microhabitat, growth under high-salinity Red Sea seawater cultivation conditions, accumulation of secondary ketocarotenoids, and increased thermal stability of extracted phycocyanin. Phylogenetic placement based on 16S rRNA phylogeny and phylogenomic analysis associated strain KAUST 171 with Halospirulina-like lineages while separating it from available Limnospira and Arthrospira genomes.

### Description

Growth was observed in BG-11 prepared with natural Red Sea seawater, with highest biomass accumulation between 35 and 40°C. Biomass concentrations reached up to 9.83 g L^-1^, with maximum C-phycocyanin content of 67.3 ± 0.77 mg gDW^-1^. Carotenoid extracts were enriched in secondary ketocarotenoids, including canthaxanthin and echinenone-like carotenoids.

### Type Material

The type strain, KAUST 171, is maintained as an active culture in the KAUST KSA Living Library.

## Acknowledgements

Kyle J. Lauersen acknowledges Baseline research funding from King Abdullah University of Science and Technology (KAUST) as well financial support provided by the Center of Excellence for Sustainable Food Security at King Abdullah University of Science and Technology (KAUST). The authors thank Professor Robert Hoehndorf and his team for conducting the Empty Quarter field expedition, enabling collection of the samples used in this study.

## CRediT author contributions

**Bárbara Bastos de Freitas**: Conceptualization, Investigation, Methodology, Data curation, Formal analysis, Visualization, Project administration, Writing - original draft, review & editing.

**Mirian dos Santos Mendes**: Investigation, Methodology, Validation. **Mark Pampuch:** Formal analysis, Software, Data curation, Visualization. **Mauricio Lopez Portillo Masson:** Investigation, Methodology, Validation.

**Kyle J. Lauersen**: Conceptualization, Resources, Supervision, Funding acquisition, Writing - review & editing.

## Data availability

The genome sequence generated in this study has been deposited in NCBI under accession number JBYDFK000000000. The data supporting the findings of this study are available within the article and its Supplementary Material. Additional data are available from the corresponding author upon reasonable request.

## Supplementary Figures captions

**Supplementary Figure S1.** Maximum likelihood phylogenomic tree showing the placement of *Halospirulina saudiensis* within *Spirulinaceae*. The tree was generated from concatenated conserved marker genes using publicly available genomes from *Spirulinaceae* and related cyanobacterial taxa. *Nostoc commune* HK-02 was used as the outgroup. Numbers at nodes indicate bootstrap support values. Scale bar indicates substitutions per site.

**Supplementary Figure S2.** Average nucleotide identity (ANI) analysis of *Halospirulina saudiensis* and related members of Spirulinaceae. Pairwise ANI values were calculated from whole-genome comparisons and visualized as a clustered heatmap. Numbers within cells indicate ANI values and dendrograms represent hierarchical clustering based on genomic similarity. *H. saudiensis* is highlighted in the matrix.

**Supplementary Figure S3.** Biomass accumulation of *Halospirulina saudiensis* under continuous CO_2_ supply compared with pH-controlled cultivation conditions. Biomass accumulation (g L^-1^) during 7 d of cultivation at 40 °Cunder pH 8, pH 9, and continuous CO_2_ supply conditions. Values represent mean ± SD of biological duplicates (n = 2), with biomass determined from 3 technical measurements per replicate. Culture appearance at d7 is shown for each cultivation condition.

**Supplementary Figure S4.** Thin-layer chromatography (TLC) of carotenoid extracts from *Halospirulina saudiensis* and *Limnospira platensis*. Carotenoid extracts were separated alongside carotenoid standards. Bands were annotated according to migration position and pigment assignment.

## References

Antecka, A., Klepacz-Smółka, A., Szeląg, R., Pietrzyk, D., Ledakowicz, S., 2022. Comparison of three methods for thermostable C-phycocyanin separation and purification. Chemical Engineering and Processing - Process Intensification 171. 10.1016/j.cep.2021.108563

Arenas Colarte, C., Balic, I., Díaz, Ó., Moreno, A.A., Amenabar, M.J., Bruna Larenas, T., Caro Fuentes, N., 2025. High-Value Bioactive Molecules Extracted from Microalgae. Microorganisms 13, 1–16. 10.3390/microorganisms13092018

Arnon, D.I., 1949. COPPER ENZYMES IN ISOLATED CHLOROPLASTS. POLYPHENOLOXIDASE IN BETA VULGARIS. Plant Physiol. 24, 1.

Barahoei, M., Kasiri, R., Feghhipour, S.E., Hatamipour, M.S., 2026. Maximizing lipid productivity in Spirulina platensis: Culture medium nitrogen interactions revealed by response surface methodology. Algal Res. 94, 104588. 10.1016/j.algal.2026.104588

Blanco-Sacristán, J., Nelson, C., Corrochano-Monsalve, M., Maestre, F.T., 2025. Biological soil crusts in the Arabian Peninsula: Ecological functions, current knowledge and research gaps. Cambridge Prisms: Drylands 2. 10.1017/dry.2025.10007

Bounnit, T., Saadaoui, I., Ghasal, G. Al, Rasheed, R., Dalgamouni, T., Jabri, H. Al, Leroy, E., Legrand, J., 2022. Assessment of novel halo- and thermotolerant desert cyanobacteria for phycobiliprotein production. Process Biochemistry 118, 425–437. 10.1016/j.procbio.2022.04.017

Breitenbach, J., Gerjets, T., Sandmann, G., 2013. Catalytic properties and reaction mechanism of the CrtO carotenoid ketolase from the cyanobacterium Synechocystis sp. PCC 6803. Arch. Biochem. Biophys. 529, 86–91. 10.1016/j.abb.2012.11.003

Chaiklahan, R., Chirasuwan, N., Bunnag, B., 2012. Stability of phycocyanin extracted from Spirulina sp.: Influence of temperature, pH and preservatives. Process Biochemistry 47, 659–664. 10.1016/j.procbio.2012.01.010

Chakdar, H., Pabbi, S., 2017. Algal Pigments for Human Health and Cosmeceuticals, Algal Green Chemistry: Recent Progress in Biotechnology. Elsevier B.V. 10.1016/B978-0-444-63784-0.00009-6

Chaumeil, P.A., Mussig, A.J., Hugenholtz, P., Parks, D.H., 2022. GTDB-Tk v2: memory friendly classification with the genome taxonomy database. Bioinformatics 38, 5315–5316. 10.1093/bioinformatics/btac672

Chun, J., Oren, A., Ventosa, A., Christensen, H., Arahal, D.R., da Costa, M.S., Rooney, A.P., Yi, H., Xu, X.W., De Meyer, S., Trujillo, M.E., 2018. Proposed minimal standards for the use of genome data for the taxonomy of prokaryotes. Int. J. Syst. Evol. Microbiol. 68, 461–466. 10.1099/ijsem.0.002516

Chuvochina, M., Gerken, J., Frentrup, M., Sandikci, Y., Goldmann, R., Freese, H.M., Göker, M., Sikorski, J., Yarza, P., Quast, C., Peplies, J., Glöckner, F.O., Reimer, L.C., 2026. SILVA in 2026: a global core biodata resource for rRNA within the DSMZ digital diversity. Nucleic Acids Res. 54, D334–D341. 10.1093/nar/gkaf1247

Demirel, Z., Sukatar, A., 2019. Purification of phycocyanin from isolated and identified hot spring cyanobacteria. Indian J. Exp. Biol. 57, 338–345.

DuBois, Michel., Gilles, K.A., Hamilton, J.K., Rebers, P.A., Smith, Fred., 1956. Colorimetric Method for Determination of Sugars and Related Substances. Anal. Chem. 28, 350–356. 10.1021/ac60111a017

Edgell, H.S., 2006. Arabian Deserts. Springer Netherlands, Dordrecht. 10.1007/1-4020-3970-0

Eren, A.M., Kiefl, E., Shaiber, A., Veseli, I., Miller, S.E., Schechter, M.S., Fink, I., Pan, J.N., Yousef, M., Fogarty, E.C., Trigodet, F., Watson, A.R., Esen, Ö.C., Moore, R.M., Clayssen, Q., Lee, M.D., Kivenson, V., Graham, E.D., Merrill, B.D., Karkman, A., Blankenberg, D., Eppley, J.M., Sjödin, A., Scott, J.J., Vázquez-Campos, X., McKay, L.J., McDaniel, E.A., Stevens, S.L.R., Anderson, R.E., Fuessel, J., Fernandez-Guerra, A., Maignien, L., Delmont, T.O., Willis, A.D., 2021. Community-led, integrated, reproducible multi-omics with anvi’o. Nat. Microbiol. 6, 3–6. 10.1038/s41564-020-00834-3

Fernández-González, B., Sandmann, G., Vioque, A., 1997. A new type of asymmetrically acting β-carotene ketolase is required for the synthesis of echinenone in the cyanobacterium Synechocystis sp. PCC 6803. Journal of Biological Chemistry 272, 9728–9733. 10.1074/jbc.272.15.9728

Freitas, B.B., Overmans, S., Medina, J.S., Hong, P.-Y., Lauersen, K.J., 2023. Biomass generation and heterologous isoprenoid milking from engineered microalgae grown in anaerobic membrane bioreactor effluent. Water Res. 229, 119486. 10.1016/j.watres.2022.119486

González-Portela, R.E., Romero-Villegas, G.I., Kapoore, R. V., Alammari, Z.M., Malibari, R.A., Shaikhi, A. Al, Al Hafedh, Y., Aljahdali, A.H., Banjar, R.E., Mhedhbi, E., Filimban, A., Padri, M., Fuentes-Grünewald, C., 2024. Cultivation of Limnospira maxima under extreme environmental conditions in Saudi Arabia: Salinity adaptation and scaling-up from laboratory culture to large-scale production. Bioresour. Technol. 406, 0–2. 10.1016/j.biortech.2024.131089

Guidi, F., Gojkovic, Z., Venuleo, M., Assuncao, P.E.C.J., Portillo, E., 2021. Long-term cultivation of a native Arthrospira platensis technical evidence for a viable production of food-grade biomass. Processes 9, 1–27.

Guillard, R.R., Ryther, J.H., 1962. Studies of marine planktonic diatoms. I. Cyclotella nana Hustedt, and Detonula confervacea (cleve) Gran. Can. J. Microbiol. 8, 229–239. 10.1139/m62-029

Hagemann, M., Marin, K., 1999. Salt-induced sucrose accumulation is mediated by sucrose-phosphate-synthase in cyanobacteria. J. Plant Physiol. 155, 424–430. 10.1016/S0176-1617(99)80126-6

Hamada, A., Hibino, T., Nakamura, T., Takabe, T., 2001. Na+/H+ antiporter from Synechocystis species PCC 6803, homologous to SOS1, contains an aspartic residue and long C-terminal tail important for the carrier activity. Plant Physiol. 125, 437–446. 10.1104/pp.125.1.437

Hoehndorf, R., 2026. Rub al-Khali Knowledge Graph: an integrated semantic resource for desert microbiome and geochemistry [WWW Document]. URL https://rubalkhali.science/

Holland, D.L., Gabbott, P.A., 1971. A micro-analytical scheme for the determination of protein, carbohydrate, lipid and rna levels in marine invertebrate larvae. Journal of the Marine Biological Association of the United Kingdom 51, 659–668. 10.1017/S0025315400015034

Hosseini, H., Bounnit, T., Saadaoui, I., 2026. Trends in Arthrospira sp. (Spirulina) Applications: A 15-Year Bibliometric Analysis and Systematic Review. Plants 15, 1– 22. 10.3390/plants15060857

Kirsch, F., Klähn, S., Hagemann, M., 2019. Salt-Regulated Accumulation of the Compatible Solutes Sucrose and Glucosylglycerol in Cyanobacteria and Its Biotechnological Potential. Front. Microbiol. 10. 10.3389/fmicb.2019.02139

Kłodawska, K., Bujas, A., Turos-Cabal, M., Żbik, P., Fu, P., Malec, P., 2019. Effect of growth temperature on biosynthesis and accumulation of carotenoids in cyanobacterium Anabaena sp. PCC 7120 under diazotrophic conditions. Microbiol. Res. 226, 34–40. 10.1016/j.micres.2019.05.003

Kobbin, K., Katibi, K.K., Norazhar, N., Mohd Nor, M.Z., Show, P.L., Basri, M.S.M., Noriznan Mokhtar, M., 2025. Integration of ultrasonication with liquid triphasic flotation system for the extraction of phycocyanin and protein from Spirulina platensis. Journal of Chemical Technology and Biotechnology. 10.1002/jctb.70090

Komárek, J., Kaštovský, J., Mareš, J., Johansen, J.R., 2014. Taxonomic classification of cyanoprokaryotes (cyanobacterial genera) 2014, using a polyphasic approach. Preslia 86, 295–335.

Krokos, G., Cerovečki, I., Papadopoulos, V.P., Zhan, P., Hendershott, M.C., Hoteit, I., 2024. Seasonal variability of Red Sea mixed layer depth: the influence of atmospheric buoyancy and momentum forcing. Front. Mar. Sci. 11, 1–17. 10.3389/fmars.2024.1342137

Kumawat, R., Kumar, A., Kundu, A., Abraham, G., Prasanna, R., Singh, A., Jaiswal, P., 2025. Factorial optimization of upstream process in Spirulina platensis for growth and carotenoid production. 3 Biotech 15, 1–18. 10.1007/s13205-025-04545-6

Letunic, I., Bork, P., 2007. Interactive Tree Of Life (iTOL): An online tool for phylogenetic tree display and annotation. Bioinformatics 23, 127–128. 10.1093/bioinformatics/btl529

Li, H., Guo, L., Chen, L., Zhang, F., Wang, W., Lam, T.K.Y., Xia, Y., Cai, Z., 2024. Machine learning-assisted optimization of food-grade spirulina cultivation in seawater-based media: From laboratory to large-scale production. J. Environ. Manage. 369, 122279. 10.1016/j.jenvman.2024.122279

Liang, Y., Kaczmarek, M.B., Kasprzak, A.K., Tang, J., Shah, M.M.R., Jin, P., Klepacz-Smółka, A., Cheng, J.J., Ledakowicz, S., Daroch, M., 2018. Thermosynechococcaceae as a source of thermostable C-phycocyanins: properties and molecular insights. Algal Res. 35, 223–235. 10.1016/j.algal.2018.08.037

Lichtenthaler, H.K., Buschmann, C., 2005. Chlorophylls and Carotenoids: Measurement and Characterization by UV-VIS Spectroscopy, Handbook of Food Analytical Chemistry. Wiley. 10.1002/0471709085

Llewellyn, C.A., Airs, R.L., Farnham, G., Greig, C., 2020. Synthesis, Regulation and Degradation of Carotenoids Under Low Level UV-B Radiation in the Filamentous Cyanobacterium Chlorogloeopsis fritschii PCC 6912. Front. Microbiol. 11, 1–13. 10.3389/fmicb.2020.00163

Lopes, D., Moreira, A.S.P., Conde, T., Ferreira, A., Oliveira, K., Conde, A., Ramos, A.A., Pereira, H., Nunes, C., Ventura, S.P.M., Coelho, N., Rodrigues, A.M.C., Rocha, H.R., Coelho, M., Gomes, A., Pintado, M., Coimbra, M.A., do Rosário Domingues, M., 2026. Seasonal biochemical fingerprints of outdoor cultivated Spirulina (Limnospira platensis) and Microchloropsis gaditana: insights into nutritional and functional shifts. Algal Res. 94, 104520. 10.1016/j.algal.2026.104520

Lowry, O.H., Rosebrough, N.J., Farr, L.A., Randall, R.J., 1951. Protein Measurement with the Folin reagent. Journal of Biological Chemistry 265–275.

Ma, G., Gao, Q., Yuan, L., Chen, Y., Cai, Z., Zhang, L., Hu, J., Wang, Y., Wu, S., Sun, Y., 2025. Spirulina (Arthrospira) cultivation in photobioreactors: From biochemistry and physiology to scale up engineering. Bioresour. Technol. 423, 132259. 10.1016/j.biortech.2025.132259

Mahanandia, S., Al-Zharani, M., Nasr, F.A., Alneghery, L.M., Behera, M., Sayyed, R., Al-Amri, I.S., Singh, L., Mastinu, A., 2025. Cyanobacterial phycocyanin: An emerging green biomolecule with antioxidant activities and potential health benefits. Int. J. Food Sci. Technol. 60. 10.1093/ijfood/vvaf196

Maneechote, W., Pathom-aree, W., Sriket, N., Wichaphian, A., Pekkoh, J., Cheirsilp, B., Srinuanpan, S., 2025. Optimizing the nutrient and light conditions for the enhancement of biomass and high value molecules productions in Spirulina using hydroponic effluent - A sustainable circular economy approach. Biomass Bioenergy 194, 107638. 10.1016/j.biombioe.2025.107638

Marin, K., Zuther, E., Kerstan, T., Kunert, A., Hagemann, M., 1998. The ggpS gene from Synechocystis sp. Strain PCC 6803 encoding glucosyl- glycerol-phosphate synthase is involved in osmolyte synthesis. J. Bacteriol. 180, 4843–4849. 10.1128/jb.180.18.4843-4849.1998

Markou, G., Kougia, E., Arapoglou, D., Chentir, I., Andreou, V., Tzovenis, I., 2023. Production of Arthrospira platensis: Effects on Growth and Biochemical Composition of Long-Term Acclimatization at Different Salinities. Bioengineering 10. 10.3390/bioengineering10020233

Marsh, J.B., Weinstein, D.B., 1966. Simple Charring Method for Determination of Lipids. J. Lipid Res. 7, 574.

Masson, M.L.P., de Freitas, B.B., Zybinskii, A., Althagafi, G., Amad, M., Fox, M.D., Lammers, P.J., Lauersen, K.J., 2025. Elevated carbon dioxide stimulates efficient organic carbon consumption for the unicellular alga Galdieria. Trends Biotechnol. 44, 547–569. 10.1016/j.tibtech.2025.10.018

Mhedhbi, E., Padri, M., Al Shaikhi, A., Al Hafedh, Y., Fuentes Grunewald, C., 2026. The potential of microalgae to contribute to sustainable animal feed production in the Arabian Peninsula, Saudi Arabia. Applied Phycology 7. 10.1080/26388081.2025.2598561

Molina-Grima, E., García-Camacho, F., Acién-Fernández, F.G., Sánchez-Mirón, A., Plouviez, M., Shene, C., Chisti, Y., 2022. Pathogens and predators impacting commercial production of microalgae and cyanobacteria. Biotechnol. Adv. 55, 107884. 10.1016/j.biotechadv.2021.107884

Nawrocki, E.P., Eddy, S.R., 2013. Infernal 1.1: 100-fold faster RNA homology searches. Bioinformatics 29, 2933–2935. 10.1093/bioinformatics/btt509

Nowicka-Krawczyk, P., Mühlsteinová, R., Hauer, T., 2019. Detailed characterization of the Arthrospira type species separating commercially grown taxa into the new genus Limnospira (Cyanobacteria). Sci. Rep. 9, 1–11. 10.1038/s41598-018-36831-0

Nübel, U., Garcia-Pichel, F., Muyzer, G., 2000. The halotolerance and phylogeny of cyanobacteria with tightly coiled trichomes (Spirulina Turpin) and the description of Halospirulina tapeticola gen. nov., sp. nov. Int. J. Syst. Evol. Microbiol. 50, 1265–1277. 10.1099/00207713-50-3-1265

Ontiveros-Palacios, N., Cooke, E., Nawrocki, E.P., Triebel, S., Marz, M., Rivas, E., Griffiths-Jones, S., Petrov, A.I., Bateman, A., Sweeney, B., 2025. Rfam 15: RNA families database in 2025. Nucleic Acids Res. 53, D258–D267. 10.1093/nar/gkae1023

Oslan, S.N.H., Xuan, N.J., Khodzori, F.A., Hakim, B.N.A., Huda, N., Julmohamad, N., Barzkar, N., 2025. Comprehensive review of bioactive compounds from microalgae as promising source for industrial applications. Algal Res. 92, 104420. 10.1016/j.algal.2025.104420

Pagels, F., Vasconcelos, V., Guedes, A.C., 2021. Carotenoids from cyanobacteria: Biotechnological potential and optimization strategies. Biomolecules 11. 10.3390/biom11050735

Palanivel, P., Thajuddin, F., Rajendran, P.J., Elumalai, A., Muralitharan, G., 2026. Exploring the phylogeny, molecular and genomic adaptations of thermophilic microalgae and cyanobacteria. Planta 263, 1–19. 10.1007/s00425-026-05002-1

Parks, D.H., Hugenholtz, P., 2026. Genome Taxonomy Database r226.0. Version 1.92. 10.15468/dpzg84

Rebelo, B.A., Farrona, S., Rita Ventura, M., Abranches, R., 2020. Canthaxanthin, a red-hot carotenoid: Applications, synthesis, and biosynthetic evolution. Plants 9, 1–18. 10.3390/plants9081039

Rippka, R., Deruelles, J., Waterbury, J.B., 1979. Generic assignments, strain histories and properties of pure cultures of cyanobacteria. J. Gen. Microbiol. 111, 1–61. 10.1099/00221287-111-1-1

Sanfilippo, J.E., Garczarek, L., Partensky, F., Kehoe, D.M., 2019. Chromatic acclimation in cyanobacteria: A diverse and widespread process for optimizing photosynthesis. Annu. Rev. Microbiol. 73, 407–433. 10.1146/annurev-micro-020518-115738

Schwengers, O., 2025. Bakta database. 10.5281/zenodo.14916843

Schwengers, O., Jelonek, L., Dieckmann, M.A., Beyvers, S., Blom, J., Goesmann, A., 2021. Bakta: Rapid and standardized annotation of bacterial genomes via alignment-free sequence identification. Microb. Genom. 7. 10.1099/MGEN.0.000685

Sinetova, M.A., Kupriyanova, E. V, Los, D.A., 2024. Spirulina/Arthrospira/Limnospira—Three Names of the Single Organism. Foods 13, 2762. 10.3390/foods13172762

Szubert, K., Toruńska-Sitarz, A., Stoń-Egiert, J., Wiglusz, M., Mazur-Marzec, H., 2021. Comparative characterization of two cyanobacteria strains of the order Spirulinales isolated from the Baltic Sea - polyphasic approach in practice. Algal Res. 55. 10.1016/j.algal.2020.102170

Tamary, E., Kiss, V., Nevo, R., Adam, Z., Bernát, G., Rexroth, S., Rögner, M., Reich, Z., 2012. Structural and functional alterations of cyanobacterial phycobilisomes induced by high-light stress. Biochim. Biophys. Acta Bioenerg. 1817, 319–327. 10.1016/j.bbabio.2011.11.008

Turland, N.J., 2025. From the Shenzhen Code to the Madrid Code: New rules and recommendations for naming algae, fungi, and plants. Am. J. Bot. 112, 1–6. 10.1002/ajb2.70026

Udayan, A., Arumugam, M., Pandey, A., 2017. Nutraceuticals From Algae and Cyanobacteria. Algal Green Chemistry: Recent Progress in Biotechnology 65–89. 10.1016/B978-0-444-63784-0.00004-7

Vasquez Guevara, C.M., Lucakova, S., Branysova, T., Ilko, V., Tobolka, A., Rajchl, A., Branyikova, I., 2025. Fresh spirulina biomass for human nutrition – safety, microbiology, storage, sensorics. J. Appl. Phycol. 37, 3595–3607. 10.1007/s10811-025-03631-9

Waditee-Sirisattha, R., Kageyama, H., 2023. Halotolerance, stress mechanisms, and circadian clock of salt-tolerant cyanobacteria. Appl. Microbiol. Biotechnol. 107, 1129–1141. 10.1007/s00253-023-12390-x

Wang, X., Xie, Y., Zhou, Z., Ruan, R., Zhou, C., Cheng, Y., 2025. Factors Influencing Phycocyanin Synthesis in Microalgae and Culture Strategies: Toward Efficient Production of Alternative Proteins. Sustainability (Switzerland) 17, 1–21. 10.3390/su17135962

Yi, L., Solanki, R., Moll, M., Vadlamani, A., De la Hoz Siegler, H., Strous, M., 2025. Toward sustainable phycocyanin production using halo-alkaliphilic cyanobacteria: from direct air capture of carbon dioxide to biorefinery. Front. Microbiol. 16, 1–9. 10.3389/fmicb.2025.1618123

