## Supplementary figures and images for "Cultivation and physiological characterization of a desert-derived *Halospirulina* isolate"

### Supplementary Figure 1

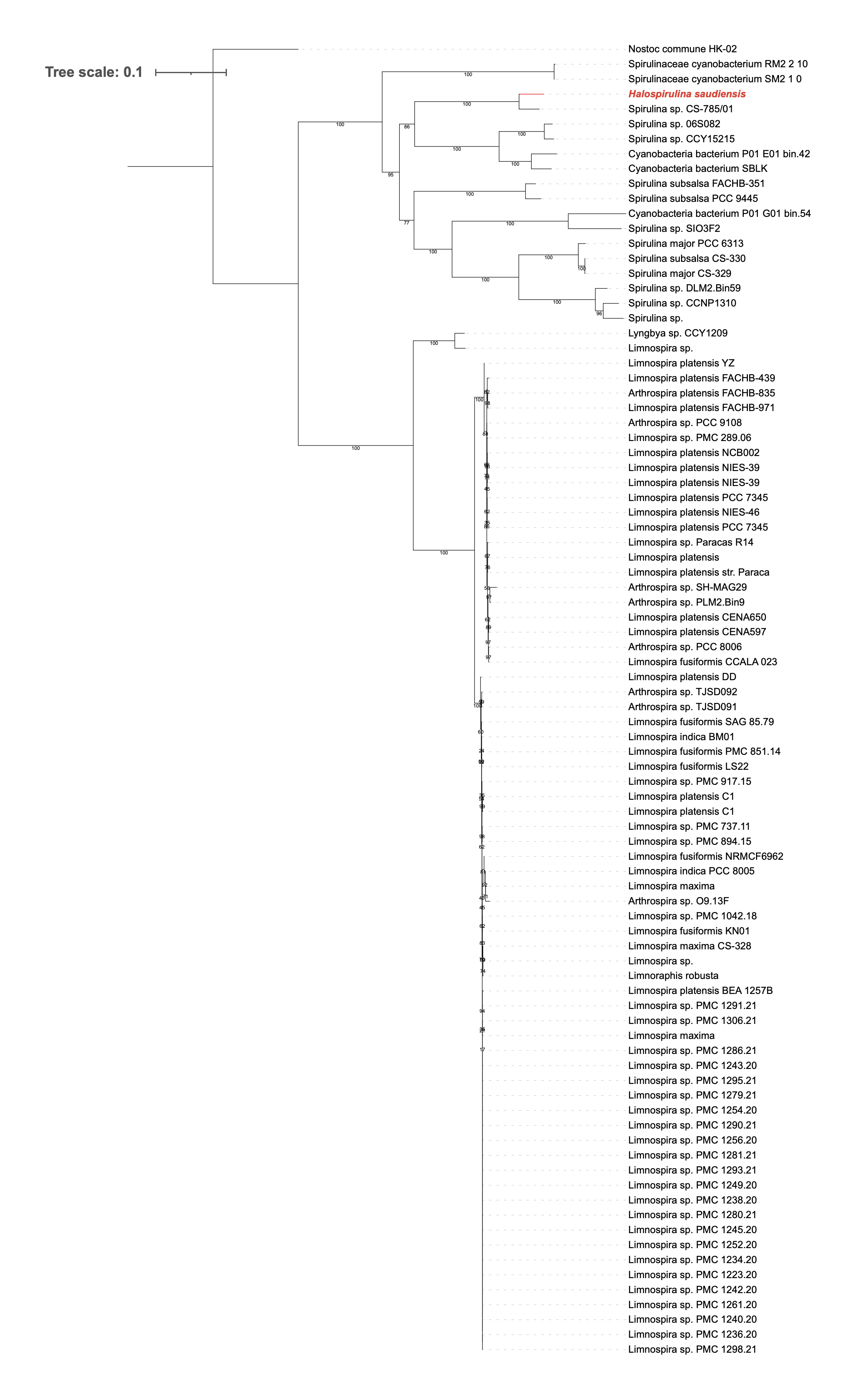

### Supplementary Figure 2

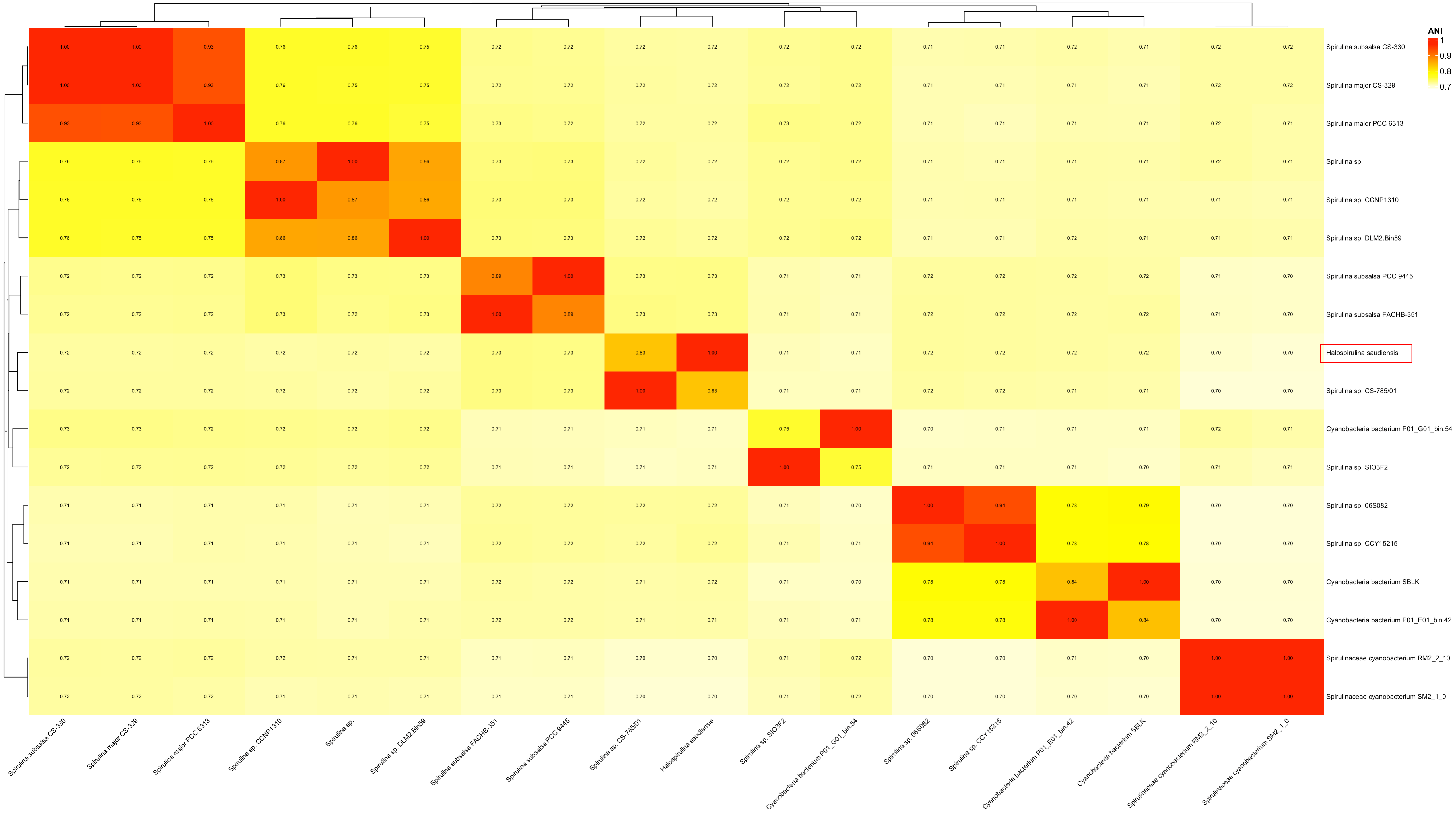

### Supplementary Figure 3

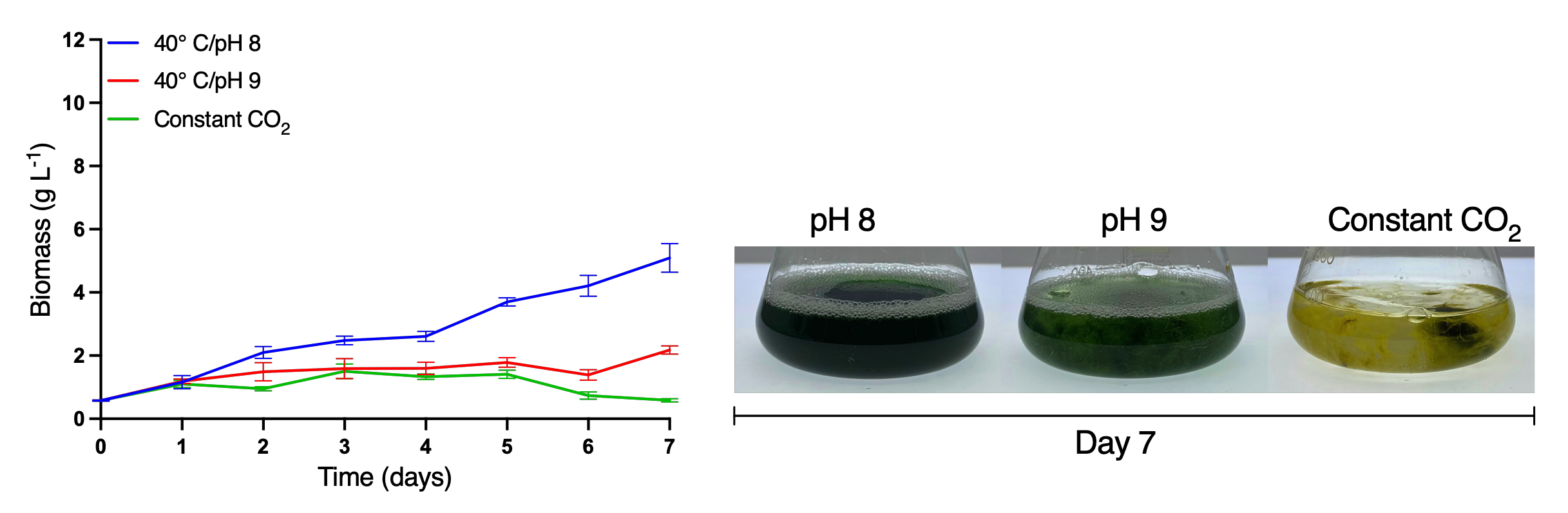

### Supplementary Figure 4

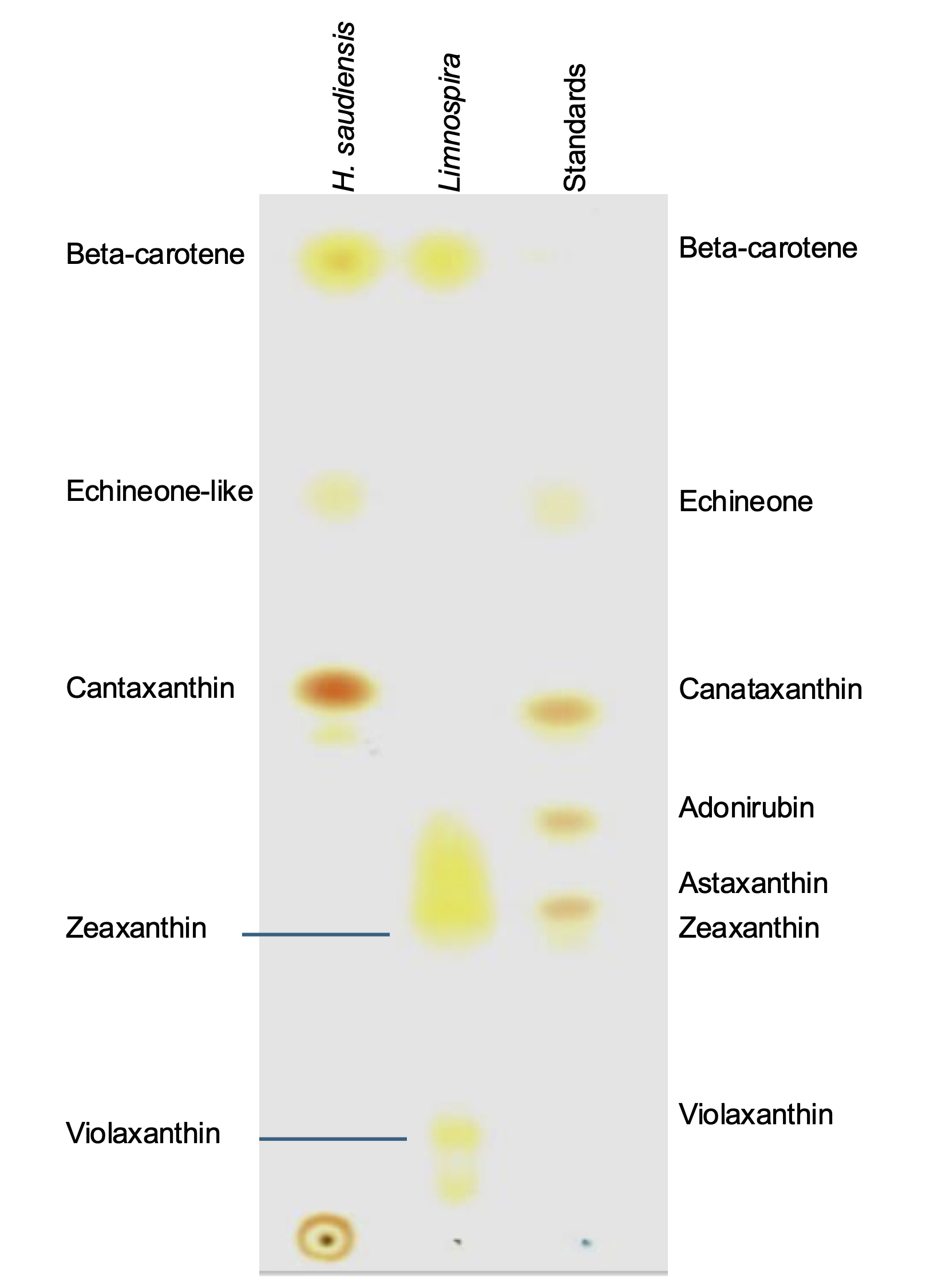
